# Shape-to-graph Mapping Method for Efficient Characterization and Classification of Complex Geometries in Biological Images

**DOI:** 10.1101/2020.03.02.972786

**Authors:** William Pilcher, Xingyu Yang, Anastasia Zhurikhina, Olga Chernaya, Yinghan Xu, Peng Qiu, Denis Tsygankov

**Affiliations:** Wallace H. Coulter Department of Biomedical Engineering, Georgia Institute of Technology and Emory University School of Medicine, Atlanta, GA, 30332, USA; School of Biological Sciences, Georgia Institute of Technology, Atlanta, GA, 30332, USA

## Abstract

With the ever-increasing quality and quantity of imaging data in biomedical research comes the demand for computational methodologies that enable efficient and reliable automated extraction of the quantitative information contained within these images. One of the challenges in providing such methodology is the need for tailoring algorithms to the specifics of the data, limiting their areas of application. Here we present a broadly applicable approach to quantification and classification of complex shapes and patterns in biological or other multi-component formations. This approach integrates the mapping of all shape boundaries within an image onto a global information-rich graph and machine learning on the multidimensional measures of the graph. We demonstrated the power of this method by (1) extracting subtle structural differences from visually indistinguishable images in our phenotype rescue experiments using the endothelial tube formations assay, (2) training the algorithm to identify biophysical parameters underlying the formation of different multicellular networks in our simulation model of collective cell behavior, and (3) analyzing the response of U2OS cell cultures to a broad array of small molecule perturbations.

**Author Summary:** In this paper, we present a methodology that is based on mapping an arbitrary set of outlines onto a complete, strictly defined structure, in which every point representing the shape becomes a terminal point of a global graph. Because this mapping preserves the whole complexity of the shape, it allows for extracting the full scope of geometric features of any scale. Importantly, an extensive set of graph-based metrics in each image makes integration with machine learning routines highly efficient even for a small data sets and provide an opportunity to backtrack the subtle morphological features responsible for the automated distinction into image classes. The resulting tool provides efficient, versatile, and robust quantification of complex shapes and patterns in experimental images.

## Introduction

Quantitative characterization of cell shapes and their organization within multicellular formations is critically important for many biomedical applications, including tissue engineering (Gupta et al. 2009), phenotypic cell-based screening (Conrad et al. 2004, Viros et al. 2008), and testing platforms for drug discovery (Murphy et al. 2010, Zanella et al. 2010). However, broadly applicable and comprehensive morphometric analysis of complex geometries in imaging data remains a challenging task. Here we present an approach that allows for an efficient and precise extraction and classification of structural features in arbitrarily complex cellular patterns, including subtle variations that are difficult to decipher using visual inspection or a set of standard geometric measures.

Currently, a number of methods have been developed for the analysis of morphological changes among individual cells (Carpenter et al. 2006, Selinummi et al. 2005, Tsygankov et al. 2014). Some targeted approaches for extracting structural features in specific applications have been also reported (Guidolin et al. 2004, Khoo et al. 2011, Lin et al. 2005, Nguyen et al. 1994), but there is still a need for a *general* methodology allowing for automated comparative analysis of complex multicellular formations. In particular, it is difficult to study the effects of a small perturbation in the extracellular environment on the collective behavior of many cells and the patterns resulting from their complex interactions (Chernaya et al. 2018). This problem is exacerbated when working with experimental systems that allow for a precise control of different physical conditions generating large and diverse sets of imaging data. To address this issue, we have developed a general approach, which automatically generates a rich set of interpretable features from images of cellular structures. These features are computed using a mathematically precise mapping of the boundaries outlining all shapes in an image onto a global graphical structure. This graphical structure captures multiple features relating to the width of the cellular objects, the shapes and roughness of the boundaries, as well as the connectivity and density of the cell clusters across the image. Using these features, we can identify images with similar structures, cluster images into groups based on structural patterns, and use the image-level characteristics for regression tasks. With this approach, one can cluster and visualize the differences between multicellular patterns based on high-level features, while still retaining the ability to interpret and understand the features defining each image type.

Unlike other graphical approaches which utilize morphological thinning (Boizeau et al. 2013, Carpentier et al. 2012, Guidolin et al. 2004) or rely on a heavily pruned skeleton (Grélard et al. 2017, Ogniewicz and Kübler 1995, Rohde et al. 2008, Styner et al. 2003, Wearne et al. 2005, Xiong et al. 2010), ours exploits the exhaustive image-scale graph to capture both fine features on the boundary of the structures and coarse features of the objects’ shapes. Furthermore, this approach is not limited to only work on networked structures. One can use this method to characterize changes in patterns of isolated cells and cell clusters, dense cellular networks, or any mixture of such formations.

As a testing system for our methodology, we first used an endothelial tube formation assay along with a computational model that simulates the formation of cellular patterns under controlled perturbations of the biomechanical properties of the cells. The tube formation assay is a useful *in-vitro* tool to screen for treatments that affect early stages of vasculogenesis. Healthy vascular endothelial cells cultured on Matrigel form dense cellular networks across the dish. Environmental or genetic perturbations can alter the resulting structure, leading to more irregular networks or completely isolated cell clusters. The standard approach to quantify these assays is to count the number of tubules (connections between cell clusters) or measure the percent coverage of a cellular network within a certain field of view (Arnaoutova and Kleinman 2010). While these approaches can be used to screen for treatments that are strongly pro- or anti-angiogenic, they are not precise enough to distinguish between more similar patterns.

For experimental perturbation of collective cell behavior, we used knockdowns of the three CCM proteins, with and without treatment by a Rho-associated protein kinase (ROCK) inhibitor, H1152 (Chernaya et al. 2018). These knockdowns all negatively affect tube formation and lead to either small isolated cell clusters or sparse patterns with large tubules depending on the targeted protein. Inhibition of ROCK partially rescues tube formation, increasing both tubule count and coverage, although the resulting cellular networks appear much more disorganized compared to wild-type. Here we show that features from the shape-to-graph mapping can differentiate images from these experiments, including the cases when images do not seem to be distinguishable and explain the differences between these visually similar groups using the features extracted from the mapping.

In addition to in-vitro assays, we utilized a simulated model that allowed us to generate a range of different multicellular patterns depending on two biomechanical characteristics: the stability of cell-cell contacts and the strength of cell-matrix adhesion (Chernaya et al. 2018). Altering these properties can create structures ranging from completely isolated cellular clusters to interconnected networks, all with varying densities. We apply our approach to predict the model parameters used to generate each *in-silico* image, demonstrating that these features can capture the trends in the way cellular structures progressively change due to the controlled modulation of the biomechanical properties of the system.

Finally, to show that our methodology is not limited to mesh-like cell formations characteristic to specific cell types, we applied it to completely different type of data from a large imaging set publicly available at the Broad Bioimage Benchmark Collection [BBBC022v1] (Gustafsdottir et al. 2013, Ljosa et al. 2012). Specifically, we analyzed confluent cultures of U2OS cells subjected to an extensive set of small molecule treatments. The global (image-scale) nature of our graph structure, which captures both the shapes of all individual cells and their relative spatial positioning in the field of view, allowed us to outperform the conventional shape metrics in terms of precision and sensitivity of the phenotypic classification.

Collectively, the performed data analysis illustrated the power of our approach for both single cell and multicellular pattern characterization, capturing apparent and subtle geometric variations using a small set of images or a large high throughput scans, while providing a way to backtrack and interpret geometric features responsible for the classification outcome.

## Results

### Shape-to-graph Mapping

Our shape-to-graph mapping is a generalization of the Voronoi Diagram to accept the edges outlining a shape as inputs. The traditional Voronoi Diagrams only operate discrete sets of points such as in the default MATLAB algorithm (Aurenhammer 1991). Our algorithm is based on a sweep-circle method (Xin et al. 2013) modified to work with line inputs. The algorithm has *O*(*n* log *n*) complexity, where *n* is the number of inputs, which scales linearly with image resolution provided the same image content. Thus, the first step in the processing pipeline is to take any binary images as an input, and output a graphical structure, which maps all piecewise linear boundaries in the image to a unique image-scale graph spanning both the foreground and background of the image **(Fig. 1)**.

**Figure 1.**
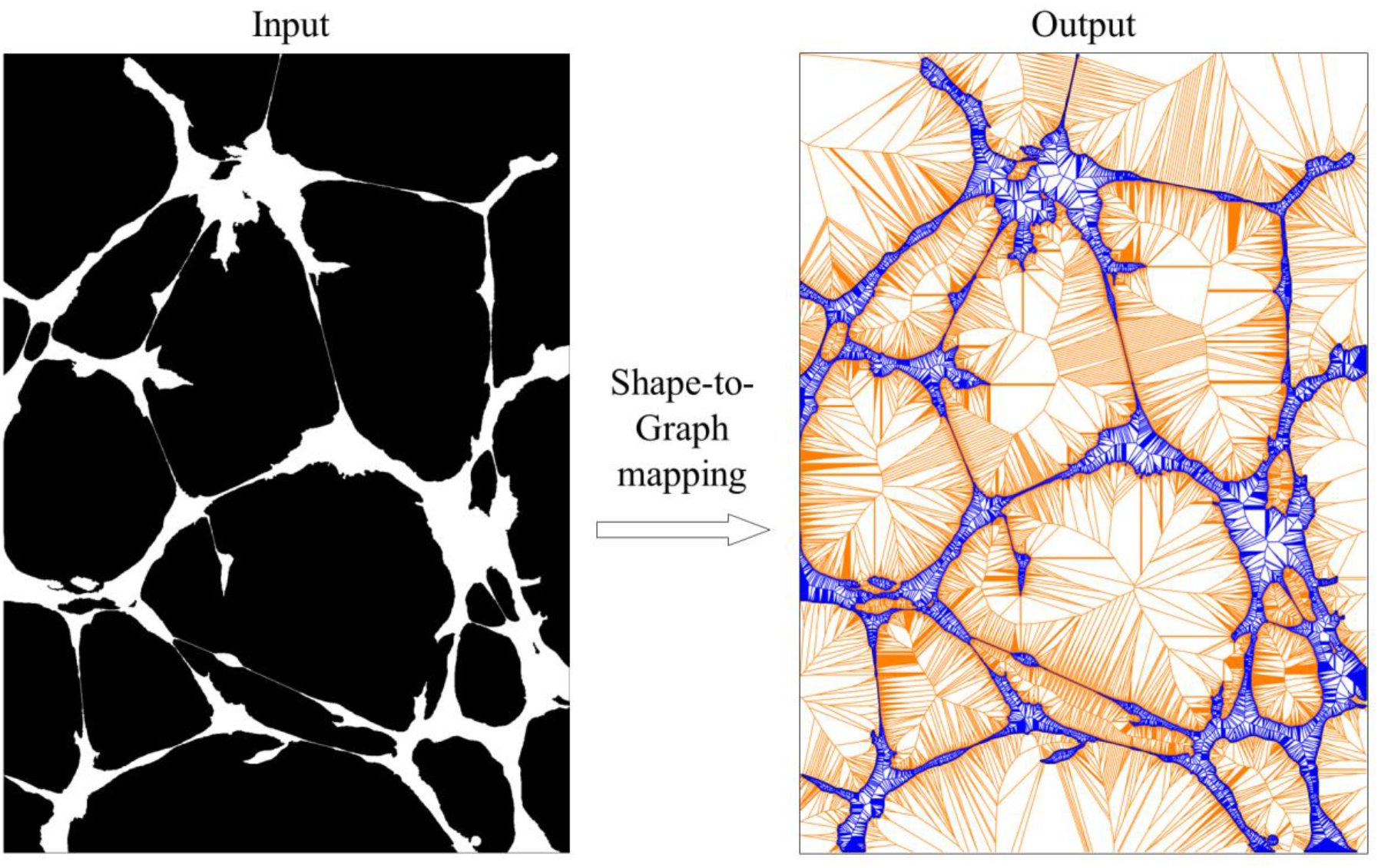
An illustration of the shape-to-graph mapping. Algorithm input is a binary image with the foreground (value 1) shown in white and the background (value 0) shown in black. Algorithm output is an image-scale graph structure. The part of the graph in the foreground (defined later in the text as in-graph) is shown in blue, while the part in the background (out-graph) is shown in orange.

### Graph construction process

A Voronoi diagram consists of vertices, which are the centers of the largest circles that can be packed within a given set of inputs, such that no input element lies within the circles. Thus each graph vertex is the center of a circle tangent to three or more input elements, while the graph edges are bisectors between two inputs. Our graph satisfies these definitions but presents a generalized version of the Voronoi Diagram, which is derived from inputs that can include both a set of points and a set of line segments. However, our main interest is an input of pixel-scale line segments forming the boundaries in a binary image.

This graph can be constructed by searching through all circles tangent to any combination of three inputs, and removing circles which contain an input within it. However, this approach would have *O*(*n*^3^) complexity, where *n* is the input size. Instead, we use a sweep-circle method, in which we compute the Voronoi diagram within an expanding circle centered at the origin. Each input generates a bisector with the sweep circle (Xin et al. 2013). Such bisector can be an ellipse for a point input or a parabola for a line input. When a new input enters the sweep circle, it’s bisector will intercept with another bisector within the sweep circle. The set of all bisector segments that are not contained within another bisector is referred to as the *beachfront* (**Fig. 2**). The interceptions between two arcs of the beachfront always lie on bisectors between the inputs, which trace out edges in the Voronoi diagram. To find the graph vertices, we only need to test inputs which have adjacent arcs on the beachfront. The ordering of arcs on the beachfront are stored within a red-black balanced binary tree (Xin et al. 2013), therefore the position of a new point within the beachfront can be found with a binary search. Thus, the complexity with this approach scales as *O*(*n* log *n*) with the number of inputs. For a more detailed, formal description of the sweep-circle algorithm, see (Xin et al. 2013). Constricted this way, each Voronoi vertex has three Voronoi edges Even in cases when the Voronoi vertex is equidistant to four or more inputs, such as the center of a regular polygon, multiple Voronoi vertices are created at the same position, each with a degree of three and a zero-length edge connecting them.

**Figure 2.**
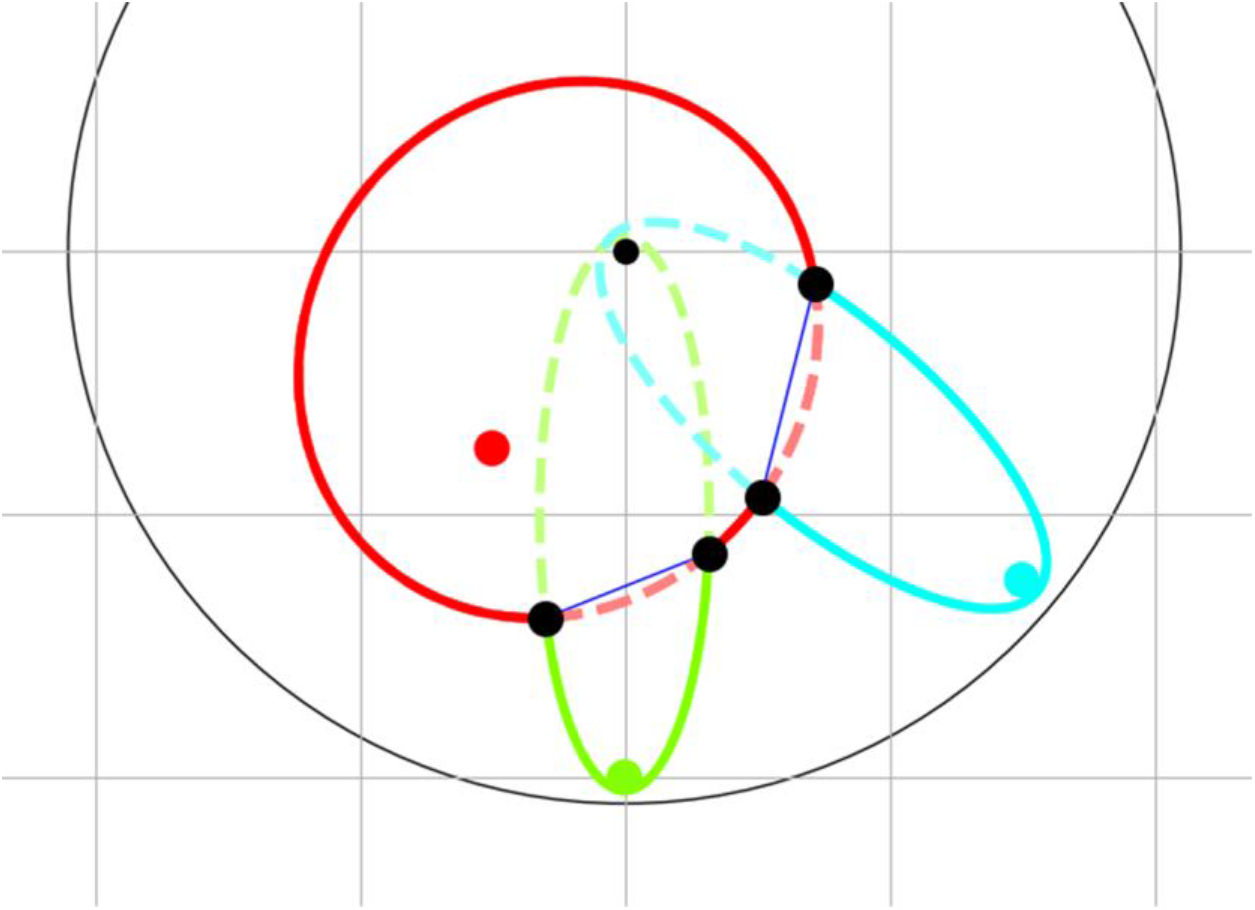
Sweep-circle Voronoi algorithm for the graph construction. In this algorithm, a sweep circle (grey circle) expands from the center of the image (grey dot). Each input point (colored dots) forms a bisector (colored ellipses) with the expanding sweep circle. The beachfront is a set of all outer most portions (solid elliptical arcs) of these bisectors. The intersections between the ellipses (black dots) trace out Voronoi edges (blue lines). When two intersection points merge, pinching out a beachfront arc, a new Voronoi vertex is formed.

To extract the algorithm input from a binary image, we trace the boundaries along the half-pixel border separating the background and foreground pixels. This is different from the conventional tracing of boundaries along the pixel centers but ensures that a horizontal or vertical line of pixels will have the width of one, rather than zero, which allows us to include pixel-size features to the image analysis. (**Fig. 3A-C**)

**Figure 3.**
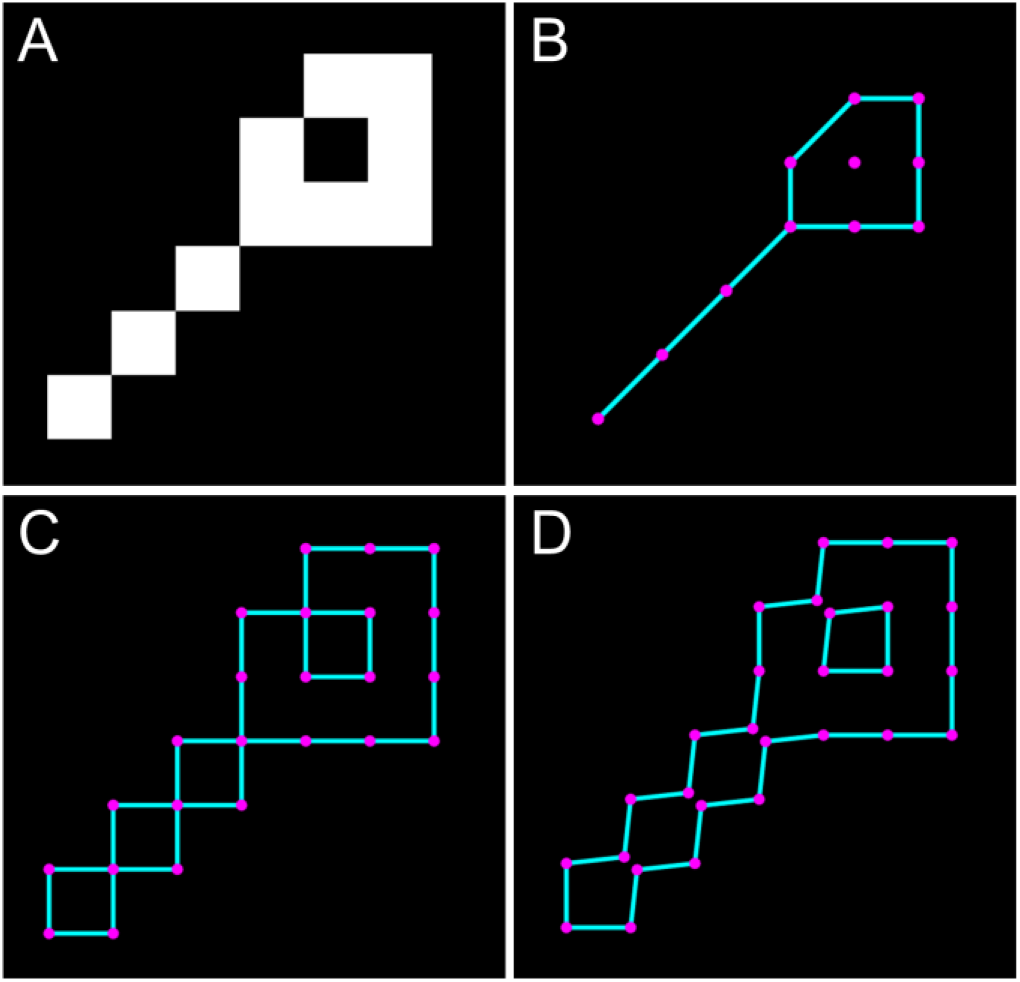
Boundary tracing. **A.** A simplified example of an input image. **B.** The conventional tracing of the boundary (implemented in MATLAB) along the centers of the pixels at the edge of a foreground object. **C.** Our algorithm traces the boundary directly along the lines separating the foreground and background pixels. **D.** An illustration of how the algorithm eliminates all boundary self-crossings by a small non-disruptive off-diagonal shift (here the shift was exaggerated for the illustration purposes).

If two boundary points overlap, such as when two foreground pixels are connected diagonally, these points are separated in the off-diagonal direction by a very small distance (we used 1/20 of the pixel size) to ensure that boundaries in the image never intersect or self-cross but unambiguously enclose the corresponding objects and holes (**Fig. 3D**).

### Graph annotation

All connected components in the foreground (objects) and background (holes) of the binary image are identified and assigned a unique numerical label. Boundaries are additionally categorized into two types: *exterior boundaries* that completely enclose a foreground object and *interior boundaries* that enclose a hole and, in turn, are enclosed by an object (**Fig. 4A**). Once the complete graph is contracted, we will refer to the part of the graph situated in the image foreground as *in-graph* and the part in the image background as *out-graph* (**Fig. 4B**).

**Figure 4.**
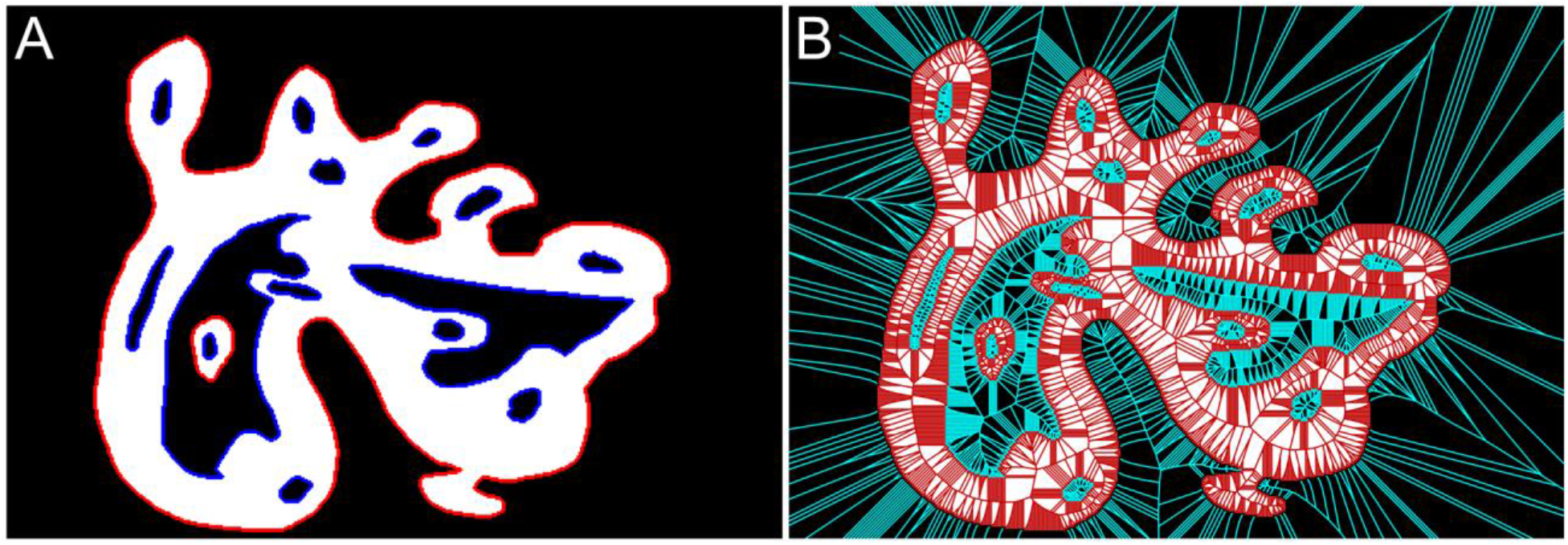
**A.** Boundary annotation: exterior boundaries are shown in red, while interior boundaries are should blue. **B.** Overall graph annotation: in-graph is shown in red, while the out-graph is shown in cyan.

Graph vertices that are equidistant to exactly two different boundaries form a sequence of vertices that we call *bridges*. Different bridges come together at graph vertices that are equidistant to three or more different boundaries and identified here as *hubs* (**Fig. 5A**). Additionally, a sequence of vertices that connect two looped bridges associated with the same boundary, which may occur when there is a hole within an extended protrusion of an object, is referred here as a *connector*. Identifying all bridges, hubs, and connectors allows us to partition the whole graph into non-overlapping *subgraphs* uniquely associated with each interior or exterior boundary (**Fig. 5B**). Extracting features from their subgraphs is central to our methodology.

**Figure 5.**
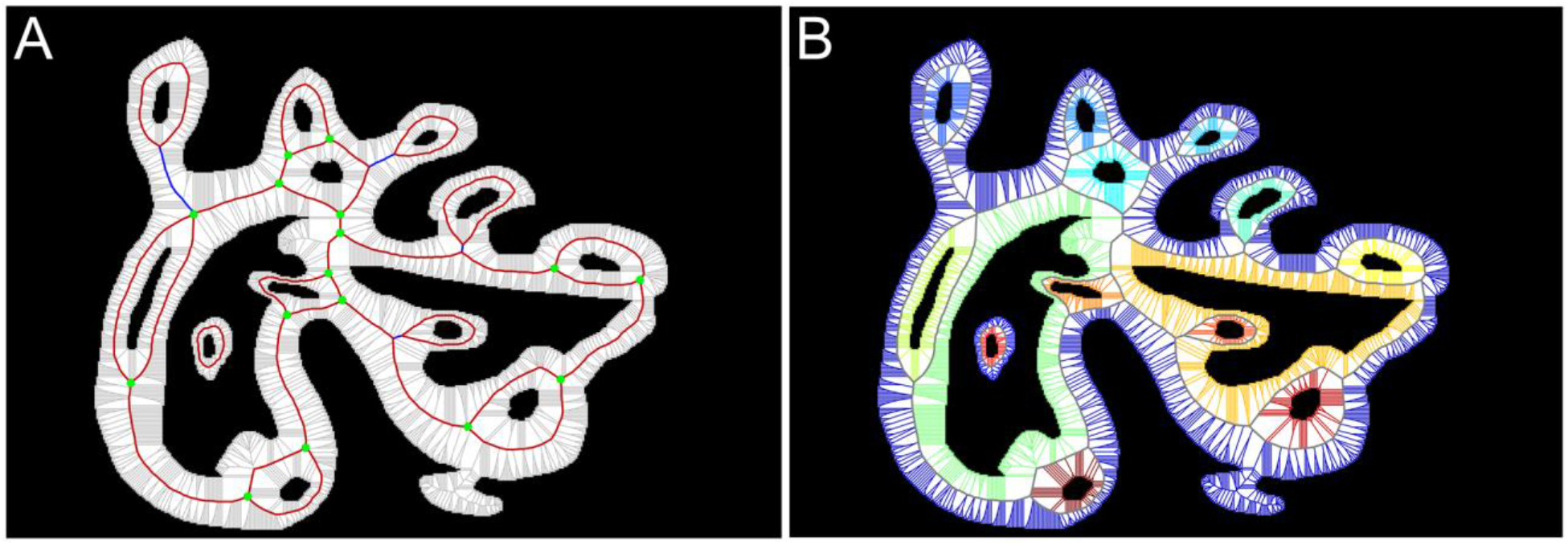
The key elements of the graph. **A.** All bridges (red), hubs (green), and connectors (blue) of the in-graph. **B.** Partitioning of the in-graph into subgraphs (shown with unique colors). Each non-overlapping subgraph is associated with exactly one interior or exterior boundary.

### Graph-based Feature Extraction

Each vertex in the constructed graph represents the center of a circle inscribed within the object. A subgraph with no bridges, such as the graph within a single-boundary object with no holes or a hole with no objects inside, is a single tree with the root node being the center of the largest inscribed circle. Otherwise, a subset of vertices located on the graph bridges and connectors of the associated subgraph acts as a set of the roots, from which graph edges branch out towards the corresponding boundary (**Fig. 6A,B**). Constructed this way, each subgraph is outlined by the boundary on one side and by a continuous sequence of bridges and connectors on the other side. We will call this sequence of bridges and connectors the *root path*. Again, in case of objects with no holes, there are no bridges, and the root path is defined as the longest path to the boundary which passes through the single root node. Based on this construction, we derive two primary metrics for each subgraph, which we call the *width profile* and the *boundary profile*.

**Figure 6.**
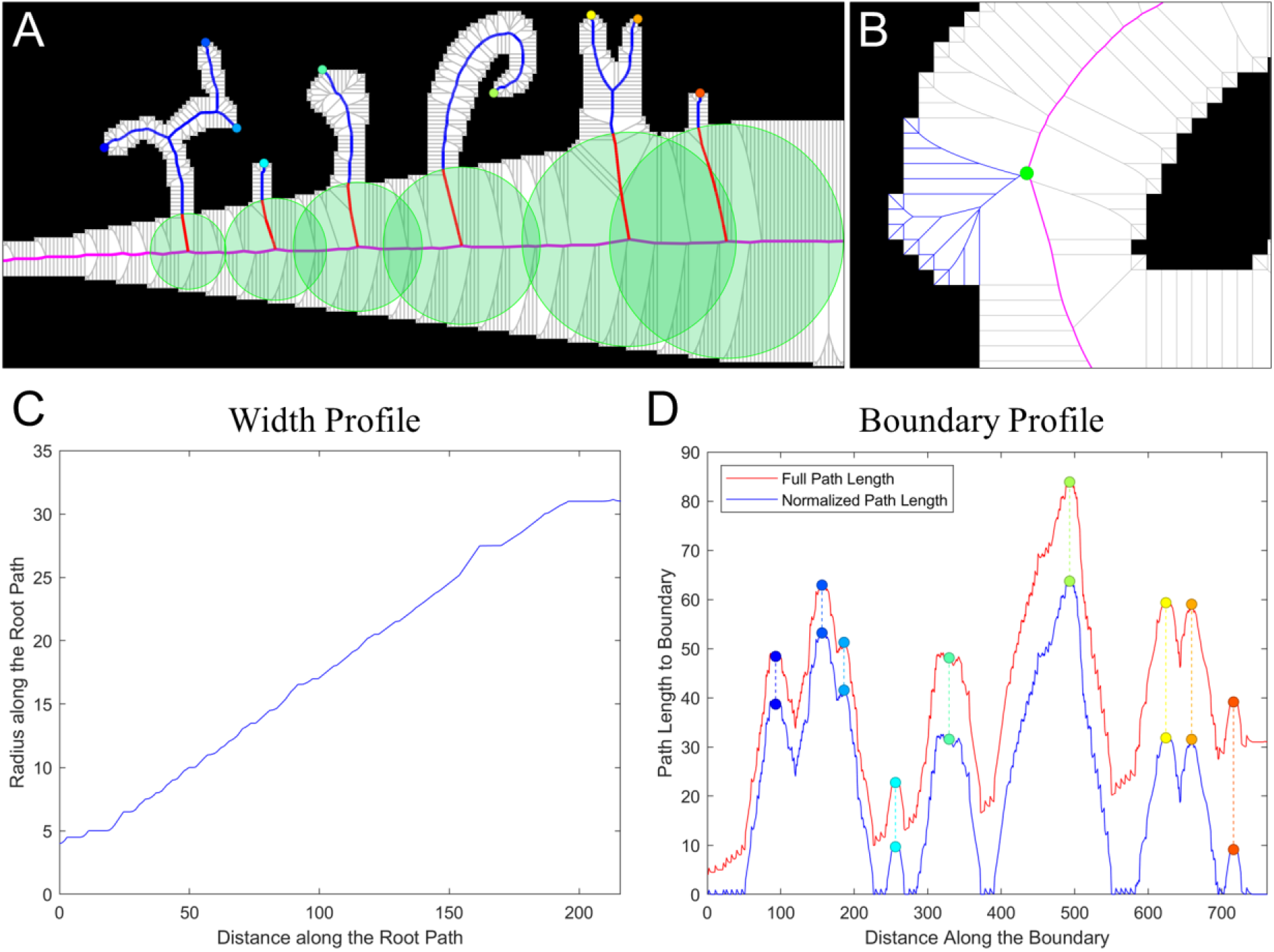
The primary graph metrics. **A.** An example of paths along the graph edges from the root path (magenta) to the tips of object protrusions. The inscribed circles (green) provide a measure for the width profile. The parts of the paths (blue) outside the circles provide a measure for the normalized boundary profile. **B.** An illustration of path (blue) branching from a root (green) to the boundary, so that each boundary point has an associated root node and a shortest path to this node along the graph edges. **C.** The resulting width profile showing the inscribed circle radii for every node on the root path. **D.** The resulting boundary profile before subtracting the radii of the corresponding root nodes (red) and after subtracting (blue). The colored points at the local maxima of the boundary profile correspond to the protrusion tips in **A.**

The *width profile* describes coarse variations in the subgraph’s width defined as the radii of the inscribed circles with the centers located at the vertices of the root path (**Fig. 6C**). When computed in background regions, this captures local variations in density. The *boundary profile* captures the size of any protrusion or bump which lies along the boundary. The boundary profile is computed by measuring the shortest distance along the subgraph edges from all points along the boundary to the corresponding root nodes. By using distances along the subgraph edges, we accurately characterize the size of these features even if the boundary is highly curved. To ensure that the boundary profile is not sensitive to the same variations in object size as the width profile, the boundary profile is normalized at each point by subtracting the radius on the inscribed circle with the center at the root node where the path to that boundary point begins (**Fig. 6D)**.

Because each boundary has a corresponding subgraph in both the in-graph and out-graph parts of the full graph, each boundary has a foreground and background width profile along with a foreground and background boundary profile. The only exception would be the most outward boundaries, for which out-graphs extend to infinity. To resolve this issue, we constrain the graph within the image by using the image boundary as the most outward boundary.

### Per-Image Structural Features

In order to characterize or compare complex geometric structures such as multicellular patterns, per-boundary classification would be insufficient as we must consider the features of all boundaries to account for the overall structure of a pattern in an image. Thus we construct a set of per-image features derived from our graph-based per-boundary features.

To this end, we start with associating each boundary with 40 features, including distribution metrics for the width profile and boundary profile, along with the area and perimeter of each boundary. Half of the features computed for each boundary come from the corresponding in-graph and half from the out-graph. The full list of features is provided in the **Supplemental Table 1**. Next, we perform k-means clustering on the list of all boundaries across all provided images (**Fig. 7**). This process creates a histogram of *N* boundary types within each image. The goal of this clustering is to automatically differentiate boundaries based on a combination of their roughness, the size and shape of the enclosed objects and holes, and the relative separation of these objects and holes. This means that holes or objects with the same shape may lie in different clusters if the cellular structure around the hole is thicker or thinner, or if the object lies in a more or less dense region. The count or frequency of the boundary types in each image then serves as a per-image feature (**Fig. 8**). The specific interpretation of each boundary type depends on the nature of data presented in the images under investigation, but this is what ultimately allows us to understand differences in the structural organization of the patterns in imaging data sets, as we show in the next section.

**Figure 7.**
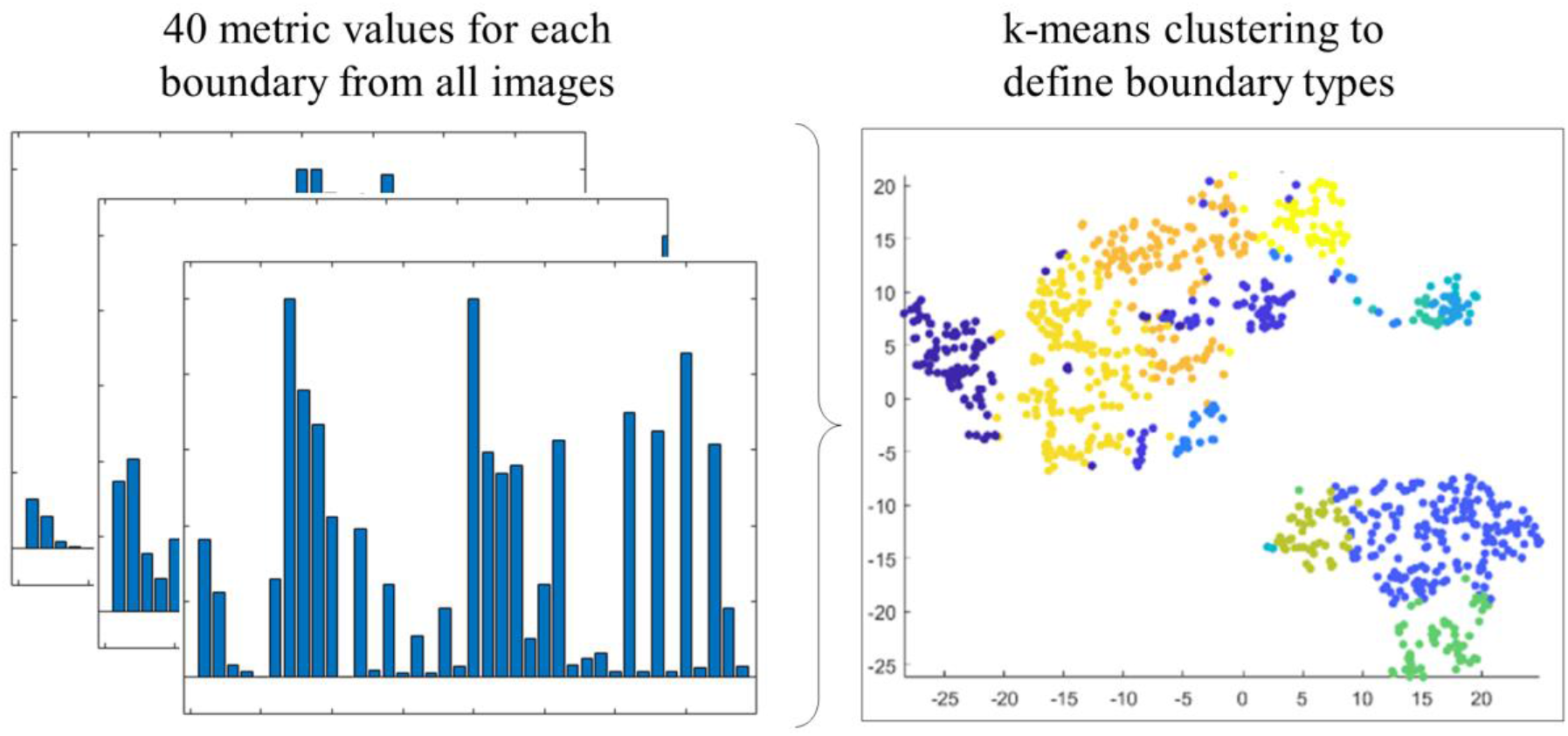
Boundary type identification. We use 40 metrics extracted for each boundary from all the images in a given set and use k-means to associate each boundary with one of the *N* classes.

**Figure 8.**
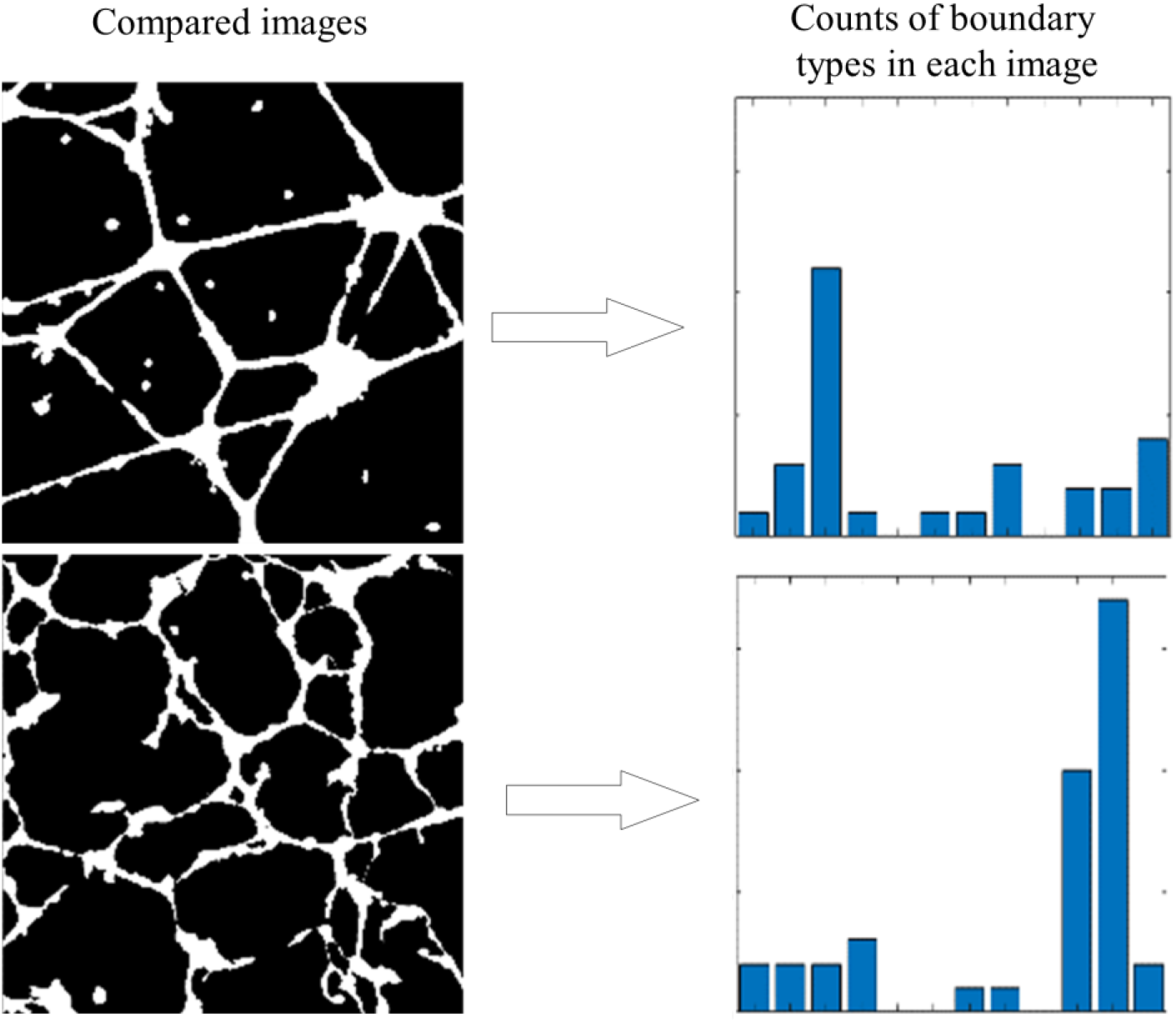
Per-image characterization. For each image, we extract the counts of boundaries that belong to each of *N* boundary types, which were determined using k-means clustering on the 40 boundary features.

### Analysis of In-Vitro Tube Formations

In this section we test the ability of our method to identify subtle structural difference in a small set of images from an in vitro endothelial tube formation assay (the experimental data has been previously published in (Chernaya et al. 2018)). The set includes images of the control cell (wild-type HUVEC) and cells with knockdown (KD) of the three Cerebral Cavernous Malformation (CCM) proteins, CCM1 (or KRIT1), CCM2, and CCM3 (or PDCD10), which disrupts the integrity of multicellular mesh. In addition, the control and KD cells were treated with an inhibitor of Rho-associated protein kinase (ROCK), which was shown to be over-activated in CCM KD cultures (Chernaya et al. 2018). The treatment with the ROCK inhibitor H1152 partially rescues the wild-type (WT) phenotype, although the resulting cellular patterns in the tube formation assay do not closely match the WT patterns. Previously, we showed that although the diseased and the H1152 treated phenotypes are clearly different from the *untreated* WT phenotype, some treated cultures are indistinguishable from the *treated* WT cells both visually and based on the traditional geometric measures(Chernaya et al. 2018). Here we show that our shape-to-graph approach allows us to identify the distinguishing features in all the phenotypes, including the ones with subtle disparities that are not apparent upon visual inspection. The latter are of the main interest from the methodology testing perspective.

For each of eight phenotypes (WT, CCM1, CCM2, CCM3, WT^H1152^, CCM1^H1152^, CCM2^H1152^, CCM3^H1152^), we used five representative fields of view **(Fig. 9A)**. The boundaries were clustered into 12 boundary types using k-means clustering. The optimal number of boundary types was selected by performing 3-nearest neighbor classification on each image, where the class of each image was determined by the class corresponding to the three most similar boundary type histograms in the image set. Twelve clusters had a 90% classification accuracy (**Supplemental Fig. S1)**.

**Figure 9.**
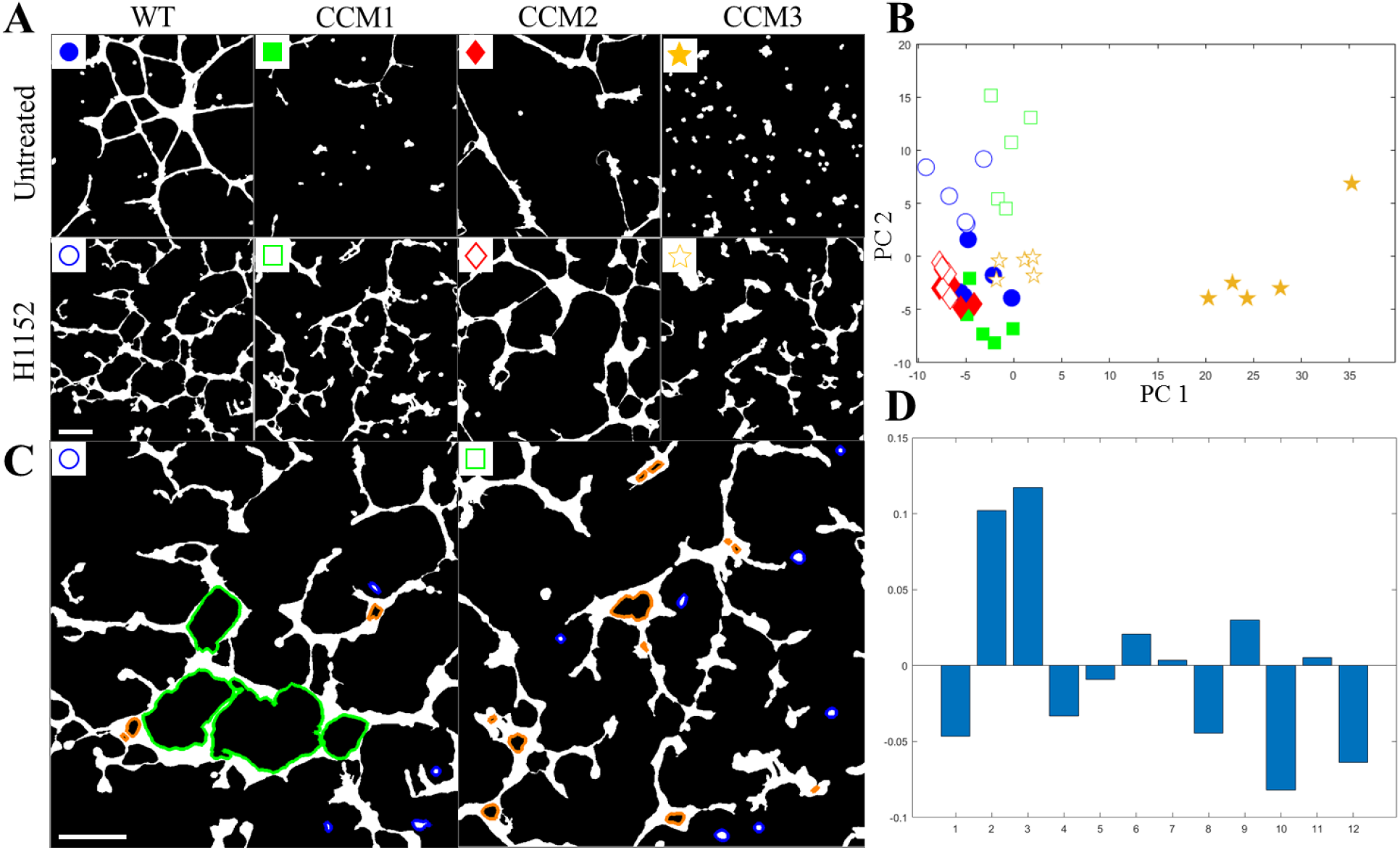
Comparison of in-vitro tube formation assay structures with eight different phenotypes. **A.** The eight phenotypes resulted from WT and the knockdown of three CCM proteins, all with and without treatment by the ROCK inhibitor. Knockdown of the CCM proteins is associated with the disruption of the otherwise connected mesh. ROCK inhibitor leads to a more connected but still noticeably disorganized network. The scale bar is 200 μm. **B.** The first two principal components of each image’s boundary type histogram. Images of a similar type and appearance tend to have similar histograms. Here, the markers indicate the corresponding images in **A. C.** Two images from WT^H1152^ and CCM1^H1152^ that appear visually similar but have significantly different boundary type counts. Boundaries that are responsible for the difference are highlighted in blue and cyan. The scale bar is 200 μm. **D.** The difference of the boundary type frequency histograms for CCM1^H1152^ and WT^H1152^. Boundary types 2 and 3 (Blue, orange) corresponding to small, isolated objects and small holes in wider locations in the network, appear significantly more often in CCM1^H1152^ formations as compared to otherwise similar WT^H1152^ structures. WT^H1152^ structures tend to have more of boundary type 10 (Green), which are medium sized holes with more bumps and protrusions extending into the hole.

Principal component analysis (PCA) was performed on the matrix of per-image boundary histograms. Generally, images of the same class group together and exist in space near images with similar structural features (**Fig. 9B**). Groups that are visually distinct, such as CCM3 cultures, which have several small cellular clusters, appear far from H1152-treated cultures with fully connected cell networks. Similarly, images with thicker structures, such as in CCM2^H1152^ cultures, appear further in principal component space from images with thinner structures, such as in CCM1^H1152^ and WT^H1152^ cultures. Visually similar structures of CCM1^H1152^ and WT^H1152^ (**Fig. 9C**), which both have many thin, disorganized connections, appear nearer to each other in principal component space. Significantly different boundary types between sets of images can be identified from the average boundary frequency histograms (**Fig. 9D)**. This difference corresponds to an increased frequency of three boundary types: type 2 consists of the *small isolated objects in regions of high density* which appear more often in CCM1^H1152^ cultures (blue boundaries in **Fig. 9C**); type 3 includes small holes in thick regions of the cellular structure, which also occur more frequently in CCM1^H1152^ (orange boundaries in **Fig. 9C**); type 10 includes medium size holes, typically with more bumps or protrusions from the cellular network extending into the hole, which occurs more frequently in WT^H1152^ samples (green boundaries in **Fig. 9C**). Descriptions of the boundary types can be determined by analyzing the distribution of the original boundary metrics within each type **(Supplemental Fig. S2)**.

### Analysis of Simulated Data

We used a previously developed computational model of endothelial tube formation (Chernaya et al. 2018) to simulate 100 images of different cellular patterns corresponding to changes in two biomechanical characteristics of cell interaction.

In this simulation model, each individual cell from a large group (hundreds to thousands) of cells sparsely distributed over the substrate surface is represented as an extendable half-ellipsoid with stochastically extending and retracting protrusions. Protrusions that extend downwards are responsible for cell-substrate interactions, while protrusions that extend sideways along the surface are responsible for cell-cell interactions. Cells form attachments when protrusions either reach deep enough into the substrate, or when it reaches another cell. Retraction of the attached protrusions leads to the cell movement, changes in cell shapes, and the buildup of the mechanical stress that can lead to the contact breakage. Ultimately, because of these cell-cell and cell-substrate interactions, the multicellular system evolves to form different patterns depending on the model parameters at the cell level. Two key parameters of interest here are the stability of cell-cell and cell-ECM adhesions. With properly selected values of the parameters, the model produces a dense cellular network closely resembling wild-type endothelial cells in our in-vitro tube formation assay. Reducing the values of each parameter leads to either a more sparse network or a number of isolated cell clusters, similar to the behavior of cell with the knockdown of CCM1 and CCM3. It is important to note here that even with a fixed set of parameters, the stochastic nature of protrusion dynamics and a random initial distribution of cells make the structures resulted in simulations vary; so that multiple patterns can be generated for the same phenotype similar to the experimental data.

As we vary the two parameters representing the stability of cell contacts, our simulations allow us to generate a sequence of cell formations with progressively changing structures (**Fig. 10A)**. Variation in the stability of cell-cell contact, the parameter *κ*_*lat*_ in the probability of contact breakage 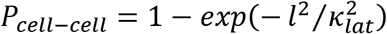, where *l* is the extension of the contact spring in the model, has a strong impact on the boundary metrics. As this parameter is increased, cells go from forming completely isolated cell clusters to a completely interconnected network. This leads to an overall reduction in boundary types corresponding to isolated cell clusters, and a shift towards networked structures with medium to large sized holes. The other parameter, *κ*_*bott*_ in the probability of cell-substrate contact breakage 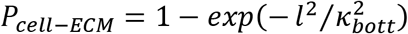, primarily affects the velocity of cell movement and the resulting density of the cell clusters. The way this parameter impacts the resulting structure depends on the network connectivity in the multicellular pattern, but generally controls the density of the structure, with low values causing cells to form larger and more sparse clusters.

**Figure 10.**
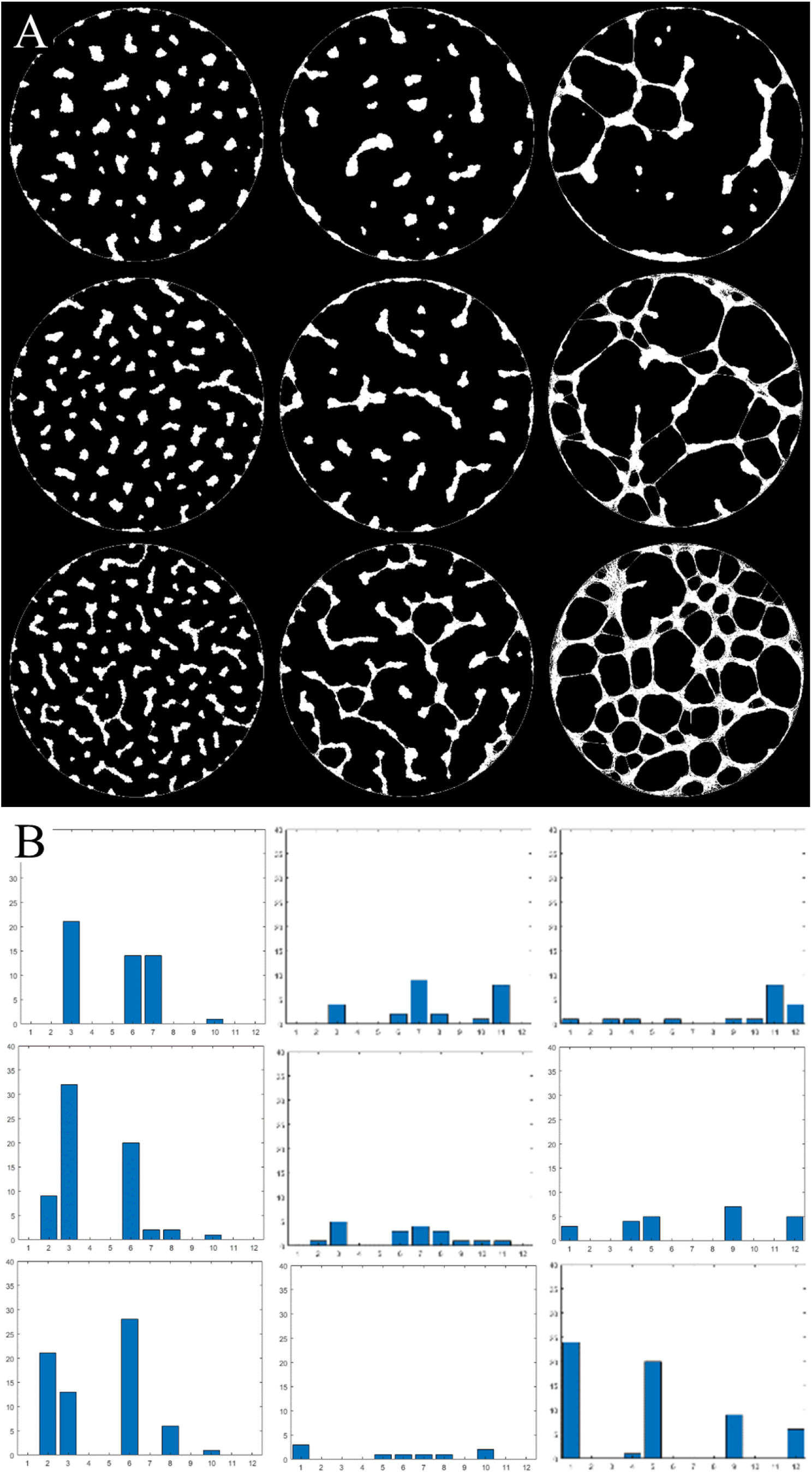
**A.** Nine representative images of multicellular formations out of 100 that were generated by varying two parameters: the strength of cell-ECM adhesion (vertical axis) and the stability of cell-cell contacts (horizontal axis). **B.** Variations of the two parameters result in visible changes in the boundary type histograms.

We applied our shape-to-graph mapping to the 100 generated images, extracted the boundary features, and clustered boundaries to create a histogram of boundary types for each image. By plotting the boundary type histograms, we can see the trends in the boundary type distribution when the two parameters are varied (**Fig. 10B**) as described above. A multi-regression model was used to predict the log-transformed values of the two model parameters based on the count of each boundary type in each image (**Fig. 11**). If these parameters have a predictable impact on the resulting multicellular pattern, and if the shape-to-graph mapping captures features that properly reflect these changes, then this multi-regression model should be able to reproduce trends in the two model parameters purely from the structural aspects of the cell patterns in the resulting images. Indeed, our approach allowed us to predict the parameter values with high accuracy: log-transformed cell-cell adhesion had a mean average error of 0.2392 with values ranging from 5 to 8 and a correlation coefficient of 0.9977, while log-transformed cell-ECM adhesion had a mean average error of 0.2782 and a correlation coefficient of 0.86695. Twelve boundary clusters were used based on cross validation performance.

**Figure 11.**
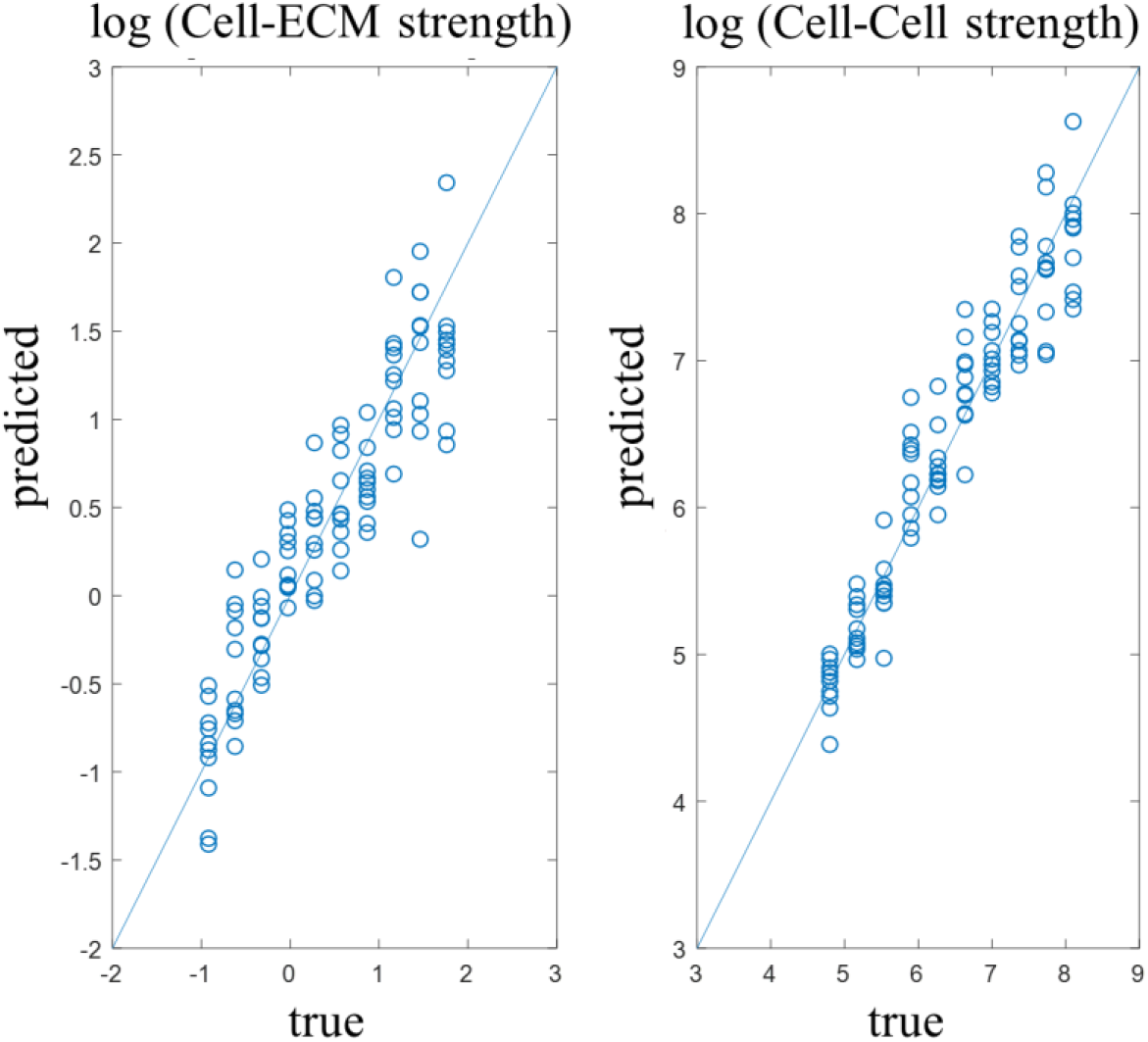
A linear regression model was trained to predict log-transformed model parameters from the boundary type histograms. The mean average error in predicting cell-cell adhesion was 0.2392, while predicting the strength of cell-ECM adhesion had the mean average error of 0.2782.

### Analysis of individual cells in a high throughput assay profiling small-molecules-induced cell cultures

In the previous sections we have focused on the analysis of complex multicellular formation with a mesh-like structures. However, our methodology is not limited to that particular type of data and can be adapted for the analysis of any images that can be segmented into the object(s) of interest and the background. To illustrate this statement, we applied our method to analyze *individually segmented* cells in a large publicly available image set with cell cultures subjected to phenotype perturbations by a variety of small molecules. We used image set BBBC022v1, available from the Broad Bioimage Benchmark Collection (Ljosa et al. 2013). The original dataset consists of fluorescent microscopy images of U2OS cells treated with one of over 1600 compounds. Five fluorescent channels were captured for each field of view. The dyes used for visualization included Hoechst 33342 (nuclei), concanavalin A (endoplasmic reticulum), SYTO 14 (nucleoli), phalloidin (actin), and WGA (Golgi complex). A CellProfiler (Carpenter et al. 2006) pipeline provided with the dataset was used to segment individual cells in each field of view via the watershed algorithm. The samples were split into 20 plates with 384 wells each. Nine fields of view were obtained for each well.

In the previous sections, our analysis relied on the input images for the shape-to-graph algorithm in the form of binary masks, in which the extracted boundaries separated the cellular structure from the background. However, in the imaging data we use here, each cell is treated as an individual object, and therefore may share a boundary with either the background or other cells. This can cause some cell boundaries to overlap **(Fig. 12A)**. To ensure cell boundaries do not overlap, we added a subpixel separation of the boundaries by shifting boundary points half-way from the previously defined half-pixel boundaries towards the corresponding pixel center **(Fig. 12B)**. This means one-pixel wide objects are thinned to have a width of half a pixel, and a half-pixel size gap is enforced to appear between two touching objects.

**Figure 12.**
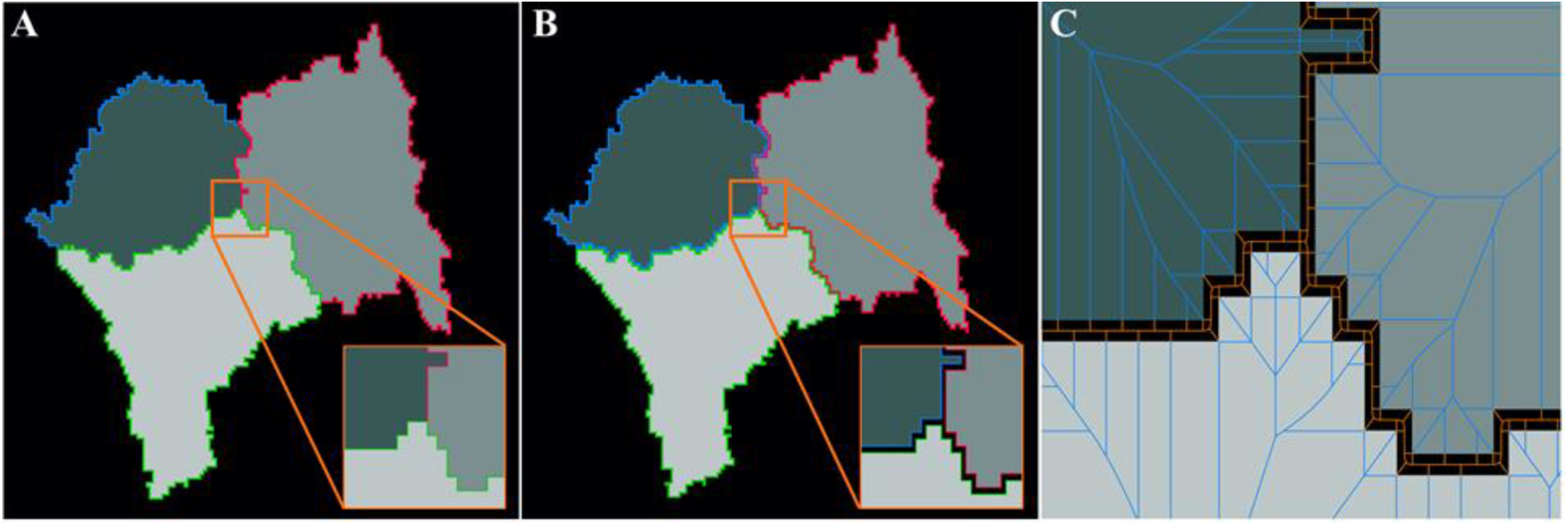
A modified boundary tracing for individual cells in a tight cluster. **A.** With the previously described boundary tracing, boundaries of contacting cells will overlap. **B.** The tracing routine is modified to place boundary points halfway between the pixel center and our original half-pixel type tracing. This creates a half-pixel gap between bordering cells. **C.** Parts of the out-graph for each cell (orange) lies within this gap. Thus, the image out-graphs will include the out-graph nodes between all the contacting cells, effectively encoding the spatial distribution of the cells in the image.

With this processing approach, the cells are presented as individual objects embedded in an image-scale mesh-like background **(Fig. 12C)**, so that the graph representation of the background (out-graph) encodes the information about the positional organization of all the cells and degree of confluency of the whole cell culture.

For our analysis, we selected 11 compounds which the authors identified as forming strong clusters based on their known mechanism of action and the 824 textural and morphological features they extracted for each image. These compound clusters include tubulin modulators (fenbendazole, oxibendazole, taxol) (**Fig. 13A)**, modulators of neuronal receptors (fluphenazine, metoclopramide, procaine) **(Fig. 13B)**, and structurally related cardenolide glycosides (digoxin, lanatoside C, peruvoside, neriifolin, digitoxin) **(Fig. 13C)**. We also included control samples from the same assays (**Fig. 13D)**. We investigated if we could predict these mechanisms of action utilizing the shape metrics derived from our shape-to-graph mapping. To this end, we extract the previously described set of measures for each object in each image. The mean and standard deviation of these per-cell metrics are computed across each well. To account for variance between plates, we subtracted the feature vector of each well by the median feature vector of the control wells in the same plate. In the end, this resulted in 208 control wells, 12 samples of tubulin modulators, 12 samples of neuronal receptor modulators, and 24 samples of structurally related cardenolides.

**Figure 13.**
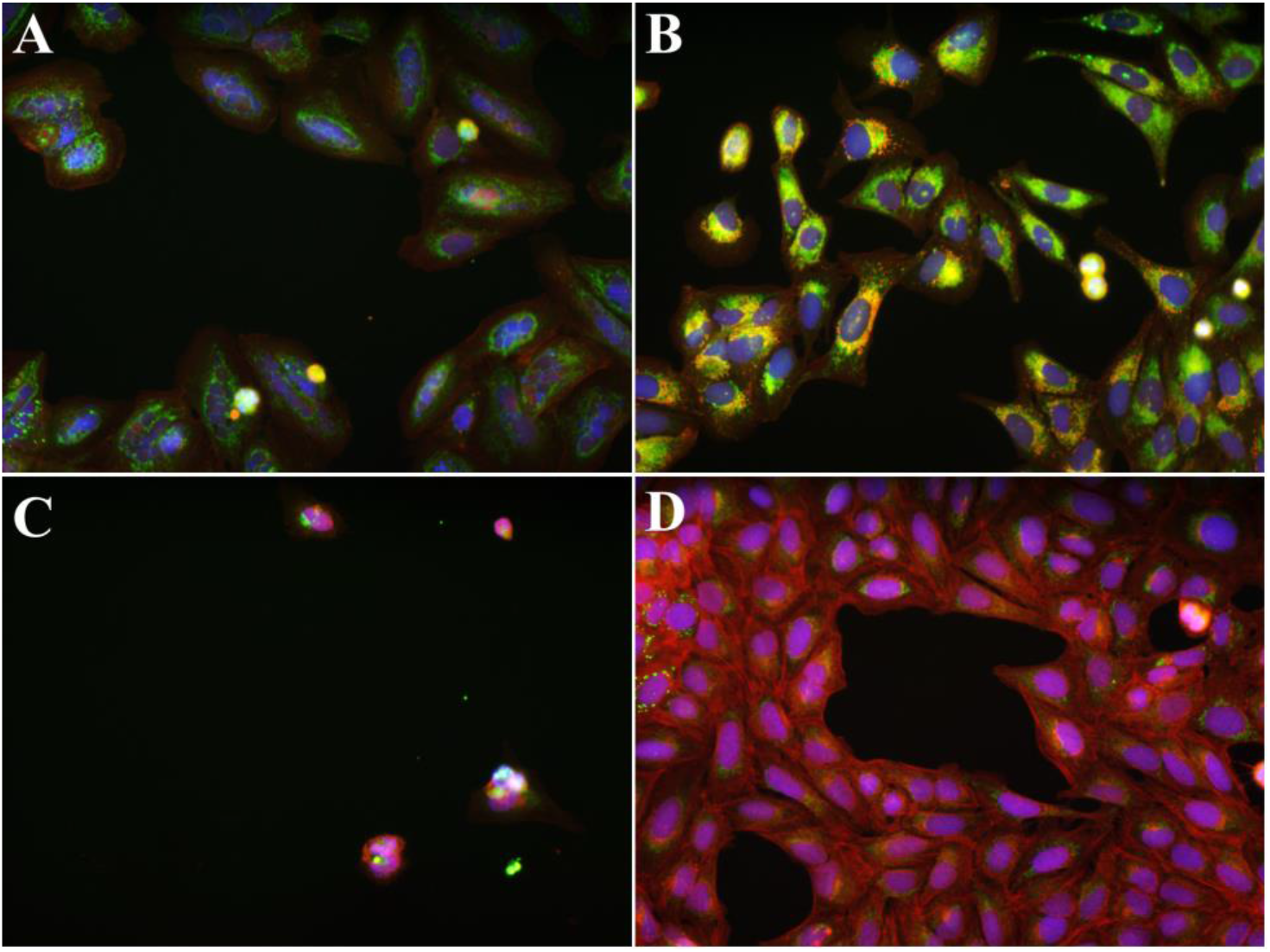
Images from the U2OS dataset. Red channel is phalloidin, blue is Hoechst 33342, and green is WGA. A) Example image from the untreated group. B) Image of cells treated with taxol from the tubulin modulators group. C) Image of cells treated with metoclopramide from the modulator of neuronal receptors group. D) Image of cells treated with digoxin from the structurally related cardenolide glycosides group.

Once the metrics were extracted, each plate was individually held-out, and a decision tree trained on the wells in the remaining 19 plates were used to predict the held-out well labels. Shape-to-graph features had a mean *F*_1_ score of 0.916 (defined as 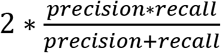, where 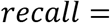 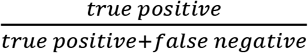, and 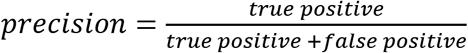), while the original published shape features (Gustafsdottir et al. 2013) had an *F*_1_ score of 0.826. Notably, the shape-to-graph mapping had much better performance on the ‘Modulators of Neuronal Receptors’ category, with a class *F*_1_ score of 0.769 versus 0.455 for the original shape features (**Fig. 14)** and each class appears to form tighter, more distinct clusters with the new features **(Supplemental Fig. S3)**. This treatment is the one which most strongly resembles the control dataset, but the cells tend to be much less dense relative to the control wells. This reduced density is captured in the out-graph radius metrics for each cell (**Supplemental Fig. S4)**.

**Figure 14.**
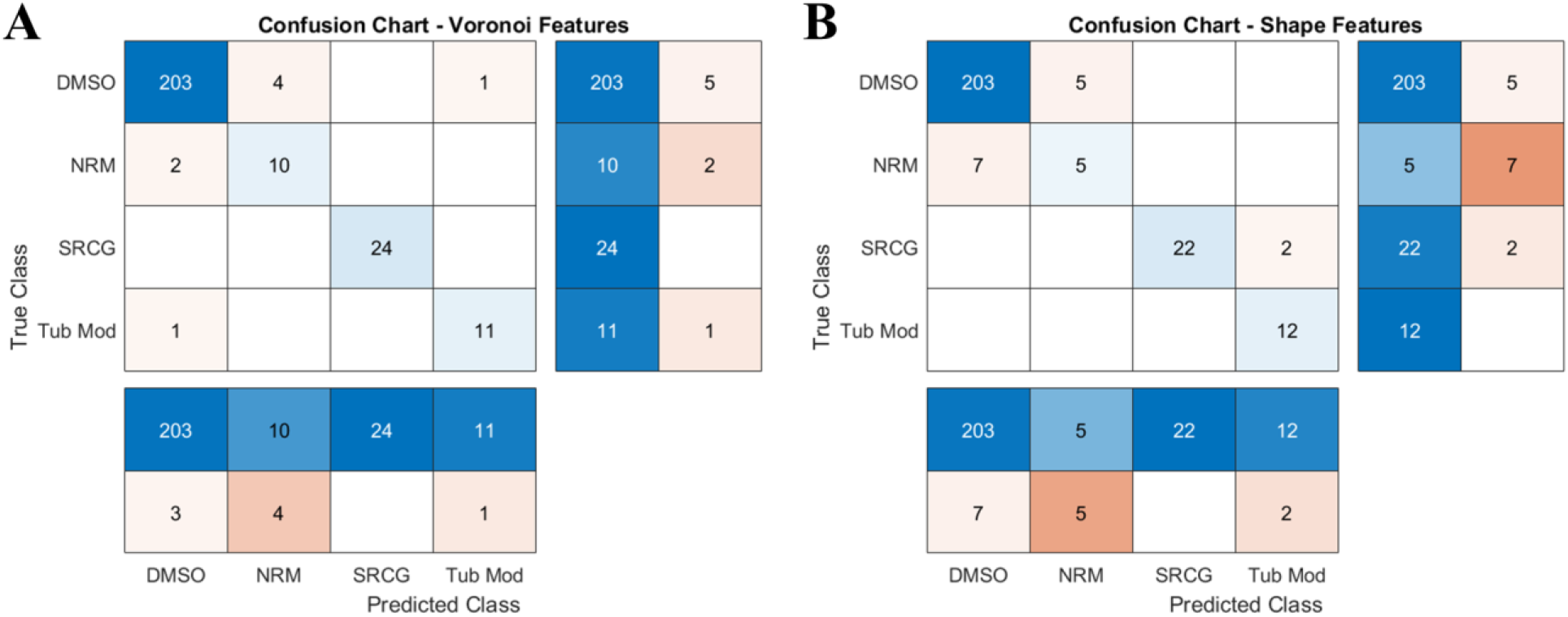
Held-out plates were classified with a decision tree trained on the remainder of the dataset. The new metrics derived with our approach tends to have better classification accuracies, especially for the control class and the modulator of neuronal receptors (NRM). Mean *F*_1_ score is 0.916 with the graph derived metrics, and 0.826 with the CellProfiler shape metrics.

### Graphical User Interface

We have created a graphical user interface (GUI) to provide readers with a quick and easy way to try our shape-to-graph mapping on their own data (**Fig. 15A**). The GUI can be used to generate and display the shape-to-graph mapping for individual images. The user can cycle through all the boundaries in the image and visualize their width and boundary profiles. A table of values of the forty measures for each boundary is also displayed.

**Figure 15.**
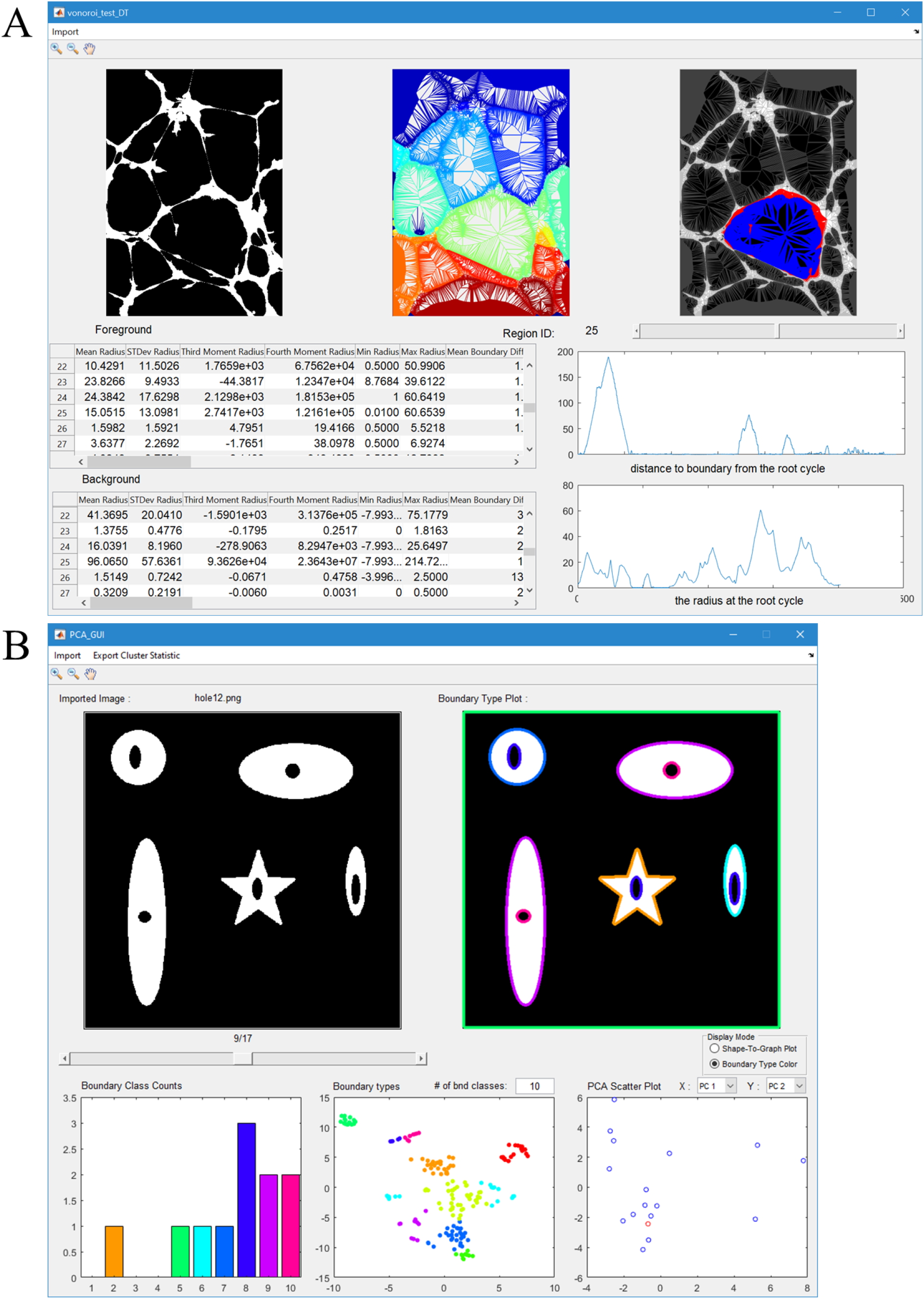
Two Graphical User Interfaces for demonstrating the graph construction and analysis. **A.** GUI for illustrating the shape-to-graph approach and the key concepts such as subgraph, in- and out-graphs, and the width and boundary profiles. The user can cycle through the boundaries and see the 40 metrics extracted for each boundary. **B.** GUI for processing multiple images. Boundaries are automatically clustered and colored according to a user-specified number of boundary types. The bottom graphs are the frequency of boundary types in the current image, a t-SNE of all the boundaries calculated by their features and colored by their resulting class, and a PCA plot of all the images derived from their boundary type histograms.

Additionally, a graphical user interface is provided to generate boundary types from multiple images **(Fig. 15B)**. The user can choose a number of boundary classes and inspect each image from the imported set with its boundaries colored according to the class they were automatically assigned based on the features from the shape-to-graph mapping (which can also be displayed). These visualizations are accompanied with (1) a color-coded histogram showing the boundary type distribution in the current image, (2) a t-SNE plot of the boundaries across all images, and (3) a plot of two user-selected principal components calculated based on the boundary type histograms across all the images. The point corresponding to the current image is highlighted in the PCA plot.

## Discussion

In this paper we introduced a methodology for extracting, quantifying, and classifying structural features of an arbitrarily complex pattern in a segmented image. The methodology is based on a mathematically defined mapping of all boundaries in the binary image onto a global graph. The graph preserves all the information specified by the boundaries but also provides an efficient and precise way of defining meaningful metrics for further processing. We illustrated the power of this approach by analyzing experimental images of human umbilical vein endothelial cells forming multicellular patterns with different levels of connectivity depending on genetic (*ccm1*, *ccm2*, *ccm3* knockdowns) and biochemical (Rho kinase inhibition) perturbations. We showed that all the visually distinguishable patterns could be reliably grouped in different classes using principal component analyses of boundary types that were defined based on a large set of graph measures. We also showed that our method is sensitive enough to identify subtle differences in visually similar patterns. More importantly, after classification, the geometric features that made such differentiation possible can be backtracked for further analysis or verification. Thus, our method allows not only for statistical quantification of pattern characteristics but also for the discovery of structural features that are not apparent from visual inspection. This is particularly important for research projects that aim to determine not only ‘which’ class of patterns a particular image belongs to, but also ‘why’ it is so in term of intuitively understandable geometric features.

As another illustration of the strength of our method, we analyzed a set of images generated with a simulation model with two control parameters responsible for the structural organization of the multicellular patterns. We showed that after training the algorithm with a subset of images, it could accurately predict the parameters used for the image generation. It is important to notice that the stochastic nature of cell-cell interactions in the model creates a variability of patterns in different simulations even with the same parameters, which can be interpreted as a noise in the data. Despite this variability, we achieved the correlation coefficients between the predicted and the actual values of the two control parameters as high as 0.9977 and 0.86695. This result shows that a biological characteristic influencing the geometry of an observed structure or pattern can be accurately quantified/predicted directly from the images once the algorithm is trained with a few images for which this characteristic was measured. One of the applications of such quantification would be an investigation of the transition dynamics between the known biological states (e.g. predicting the onset of a diseased phenotype).

Our methodology works for any binary images. Because we construct the graph for both foreground and background, the extracted features characterize the geometry of individual objects, connectivity in networked structures, as well as the relative organization of isolated objects. This fact makes our method highly versatile and generally applicable. We illustrated this statement reanalyzing a subset of previously published data set from a high throughput assay profiling small-molecule-induced U2OS cell cultures (Gustafsdottir et al. 2013). We used the same processing pipeline as in the original study but apply the geometric features from our shape-to-graph mapping. By comparing a combined metric of precision and sensitivity, the *F*_1_ score, we showed that our graph representation of the image content provides an improvement in classification performance of 10% for the three major mechanisms-of-action clusters and 40% for the cluster that differs the least from the wild type cultures. Saying that, it is important to notice that the initial, pre-processing step of segmentation is critical and the presented method can be only as accurate as allowed by the quality of microscopy and the segmentation routine.

## Materials and Methods

### Cell culture

Human umbilical cord endothelial cells HUVEC (Lonza, Walkersville, MD) were maintained in EGM-2 medium (Lonza) at 37°C/5% CO2 and passaged every 3 to 4 days for up to 6 passages at a 1:5 sub-culturing ratio. For tube formation experiments, 4.5−5×10^3^ cells were plated into each well of angiogenesis μ-slides (ibidi, Fitchburg, WI) coated with 10 μl of growth factor reduced phenol red-free Matrigel (Corning, Corning, NY), and incubated for up to 18 hrs.

### Microscopy

For endothelial tubule formation imaging, cells plated on Matrigel were incubated with CellMask™ Green Plasma Membrane Stain (Invitrogen, Carlsbad, CA) for 15 min at 37°C. The media was changed to phenol-free EGM-2 supplemented with 2% FBS and growth factors (PromoCell GmbH). Images were acquired using PerkinElmer UltraVIEW VoX spinning disk confocal microscope (PerkinElmer, Waltham, MA). Image processing and analysis were performed using ImageJ software (NIH). Images in Figure 9 represent a 1.2 mm by 1.2 mm areas. With the plating density of ~ 400 cells per mm^2^, there is ~600 cells in each image.

### Gene expression knockdown

To achieve knockdown of CCM protein expression, cells were infected with PLKO.1 vector based lentiviruses carrying shRNAs for human krit1 (RHS4533-EG889), ccm2 (RMM4534-EG216527), and pdcd10 (RHS4533-EG11235) genes (Dharmacon, Lafayette, CO). Lentiviral particles, prepared and purified by VectorBuilder technical service group (VectorBuilder, Santa Clara, CA) were added to EGM-2 media supplemented with 8μ/mL polybrene for 48 hrs. Transduced cells were selected through their resistance to puromycin added to the growth media in the concentration of 2.5 μg/ml. Expression knockdown was measured by real-time PCR with TaqMan gene expression assays. Phenotypic experiments were conducted between 6 and 10 days after infection.

### Image Preprocessing

Simulated images in vector format were rendered at 1024×1024 resolution. By design, the model generates binary images with all interacting cells and their protrusions being the foreground of the image. All holes smaller than 100 pixels were automatically filled. Multiple fields of view were sampled from experimental images of tube formation at a fixed resolution of 690×690 pixels. The images were segmented with a simple threshold followed by manual corrections to under segmented tubules. Cellular debris below 50 pixels in size were automatically removed.

Boundaries were extracted from each binary image. Linear pixel-size segments that connect boundary points serve as the input to the shape-to-graph mapping algorithm. Rather than defining boundary points at the center of each pixel at the edge of an object, points on the boundary were placed on the half-pixel border between an object and the background. This ensures that any object within the boundary has a non-zero area and any protruding part of an object has a non-zero width. When operating on label images, boundaries are extracted from the largest four-connected components for each label. Boundary points are placed half-way between the center of the pixel and the half-pixel edge used for binary images. This creates a half-pixel sized gap between objects which share a boundary, and any objects which are one pixel wide will have a width in the Voronoi diagram of 0.5px.

## Acknowledgements

We would like to acknowledge the core facilities at the Parker H. Petit Institute for Bioengineering and Bioscience at the Georgia Institute of Technology for the use of their shared equipment, services and expertise. This work was supported by the National Science Foundation grant CCF-1552784 and the ISAC Marylou Ingram Scholarship to P.Q. and by the U.S. Army Research Office (ARO) grant W911NF-17-1-0395 to D.T. and by funds from the Marcus Foundation, The Georgia Research Alliance, and the Georgia Tech Foundation through their support of the Marcus Center for Therapeutic Cell Characterization and Manufacturing (MC3M) at Georgia Tech.

## Supporting information

1. Supplemental Information with Figures and Tables in a single PDF file.
2. All Scripts and GUIs in a single ZIP file.

